# Interactive single cell RNA-Seq analysis with the Single Cell Toolkit (SCTK)

**DOI:** 10.1101/329755

**Authors:** David F. Jenkins, Tyler Faits, Emma Briars, Sebastian Carrasco Pro, Steve Cunningham, Joshua D. Campbell, Masanao Yajima, W. Evan Johnson

**Affiliations:** Division of Computational Biomedicine, Boston University School of Medicine, Boston, MA; Bioinformatics Program, Boston University, Boston, MA; Department of Mathematics & Statistics, Boston University, Boston, MA; Department of Biostatistics, Boston University School of Public Health, Boston, MA

## Abstract

Single cell RNA-sequencing (scRNA-Seq) allows researchers to profile transcriptional activity in individual cells. However, the complex nature of these data and variability in study design and data generation requires sophisticated computational tools and informed analytical decisions. Here, we present the Single Cell Toolkit (SCTK), an interactive scRNA-Seq analysis package that enables users to perform scRNA-Seq analysis interactively using a command-line workflow or a graphical user interface (GUI) written in R/Shiny.

## Main Text

Single cell RNA-sequencing (scRNA-Seq) techniques allow researchers to explore the transcriptional landscape of a sample at the resolution of the individual cell. In the context of cancer, scRNA-Seq can identify the subclonality of a tumor sample to improve our ability to identify the cell-specific mechanisms that drive tumor growth and can characterize different cellular populations within the tumor microenvironment such as immune cells^1,2^. However, different optimizations of parameters and algorithms are required for filtration, normalization, clustering, and differential expression of scRNA-Seq data compared to bulk RNA-seq due to the low amount of starting material and technical bias introduced in the common scRNA-Seq library preparation techniques^3^. Tools for normalization and analysis of scRNA-Seq data exist to overcome these technical biases, but these tools are not integrated and require command line processing of samples and knowledge of the many options available for each tool, which makes them difficult to use, especially for scientists without training in bioinformatics^4–9^. Even for more advanced users, there is still a need to interactively explore scRNA-Seq results during processing to help make dataset specific decisions that can affect downstream analysis.

Here, we present the Single Cell Toolkit (SCTK), an R/Shiny^10^ based package for both command line and interactive scRNA-Seq processing. Users can upload count and annotation data and interactively explore and perform analyses. Data and results can be saved in a convenient object for downstream command line analysis, or to reload into the GUI in another session. With the SCTK, it is possible to perform a full analysis workflow from uploading raw data to downloading processed results. While other tools can perform specific scRNA-Seq analysis steps, the SCTK is the first fully interactive scRNA-Seq analysis workflow available within the R language (**Table 1**).

**Table 1.** Comparison of SCTK and other popular scRNA-Seq analysis tools. The SCTK supports a full interactive scRNA-Seq analysis workflow and supports the SingleCellExperiment object for data storage.

| Package | SCATER<sup>4</sup> | SC3<sup>5</sup> | SEURAT<sup>6</sup> | SCDE<sup>7</sup> | PAGODA<sup>8</sup> | MONOCLE<sup>9</sup> | SCTK |
| --- | --- | --- | --- | --- | --- | --- | --- |
| Filtering and Data Summary | ✓ |  | ✓ |  |  |  | ✓ |
| Dimensionality Reduction | ✓ |  | ✓ |  |  | ✓ | ✓ |
| Clustering |  | ✓ | ✓ |  | ✓ | ✓ | ✓ |
| Batch Correction |  |  | ✓ | ✓ |  | ✓ | ✓ |
| Differential Expression |  | ✓ | ✓ | ✓ |  | ✓ | ✓ |
| Pathway Enrichment |  |  | ✓ |  | ✓ |  | ✓ |
| Power Calculation |  |  |  |  |  |  | ✓ |
| GUI | ✓ | ✓ |  |  | ✓ |  | ✓ |
| SingleCellExperiment Support | ✓ | ✓ |  |  |  |  | ✓ |

The SCTK is organized into several analysis modules (**Fig. 1**). Analysis modules include data summary and filtering, dimensionality reduction, clustering, batch correction, differential expression, pathway activity analysis, and power calculations to evaluate the tradeoff between sample size, cell numbers, and sequencing depths. All analysis modules can be run interactively through the Shiny web interface or through the R console. Modules can be run in isolation or sequentially to perform a full scRNA-Seq analysis pipeline.

**Figure 1.**
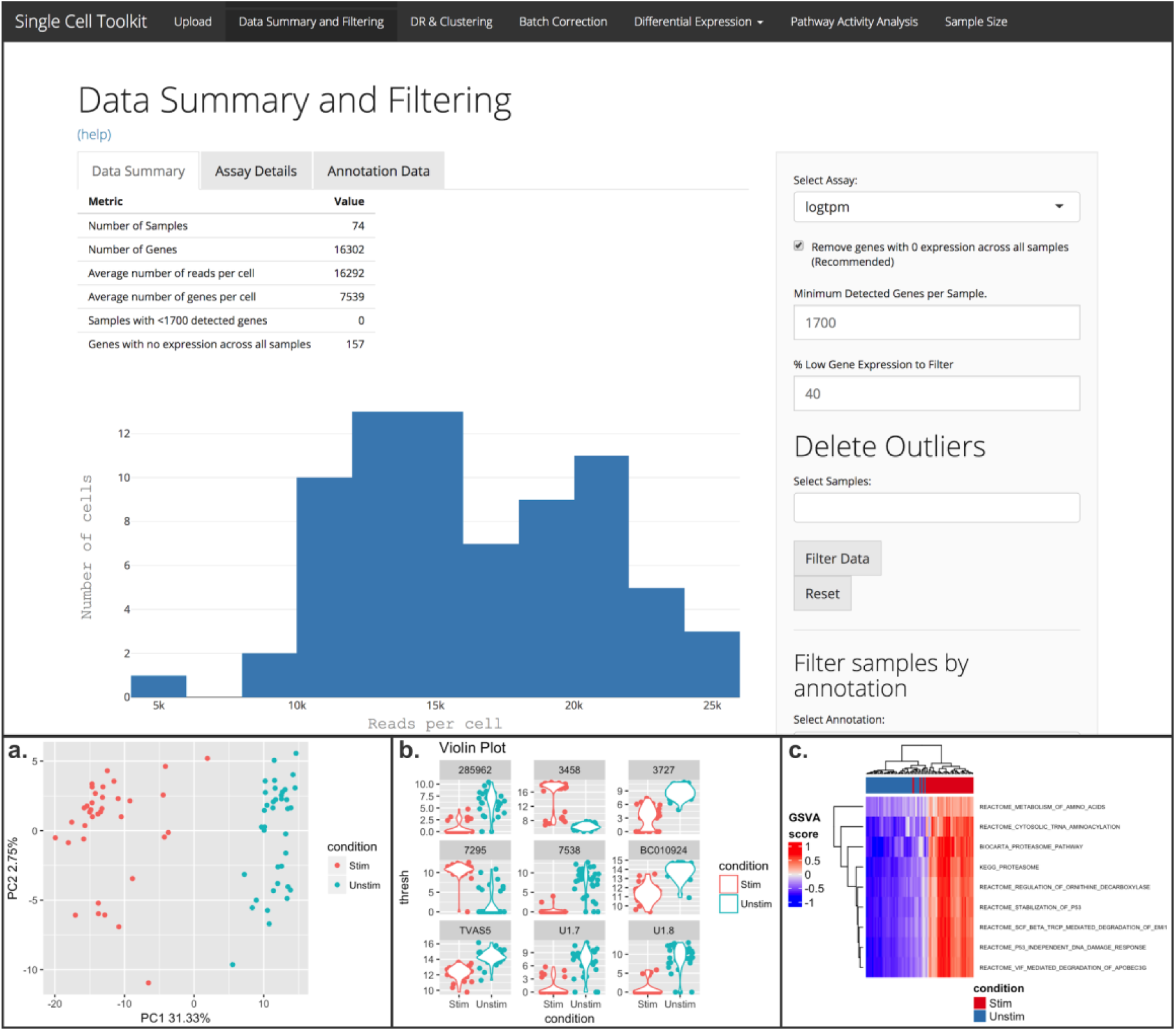
Interactive analysis modules and example output plots available in the SCTK. Top: data summary and filtering tab. **a.** PCA visualization. **b.** Violin plots of differential expression using MAST. **c.** Pathway activity analysis

Steps in the analysis pipeline are performed on a SCTKExperiment object, an extension of the SingleCellExperiment and RangedSummarizedExperiment objects developed by the Bioconductor project^11^. This object is organized into identically sized matrices designed to store counts, normalized counts, or batch corrected data; a data frame for sample annotation information; and a data frame for feature annotation information. Additionally, data from dimensionality reduction approaches such as principal component analysis (PCA) and t-distributed stochastic neighbor embedding (t-SNE) can be stored in the object. The SCTKExperiment object also stores the percent variation explained by each principal component and will continue to be expanded in the future to store additional single cell data, annotations, and results. The SingleCellExperiment object has been optimized to store large datasets by using sparse matrices and an efficient API to support data that would otherwise not fit into memory using a standard matrix^12^. The SingleCellExperiment can also store information about spike-in transcripts and sample specific size factors for normalization. By utilizing an object that can efficiently store both raw data and downstream analysis results, analysis can be performed within the SCTK, saved into the object, and loaded into R for additional analysis on the command line.

In addition to interactive analysis, experienced computational users can use the SCTK as part of an automatic pipeline. For larger datasets, such as the 10x Chromium Single Cell Solution and other droplet based high throughput methods, long running analysis steps such as t-SNE can be saved into the SCTKExperiment object and be brought into the SCTK for visualization or further interactive analysis^13^.

We applied the SCTK and our workflow on multiple data examples, including stimulated and unstimulated mucosal-associated invariant T cells, and induced pluripotent stem cells from Yoruba male reference samples to identify batch effects^14,15^. These example datasets show the SCTK’s ability to identify biologically meaningful results through R console analysis using SCTK R functions, or through the GUI (**Supplementary Note 1**).

Additionally, we have used the analysis workflow available in the SCTK to identify pathway activity differences between breast cancer cells before and after drug treatment, indicating that the SCTK can be used to identify biologically meaningful results. By performing pathway activity analysis on this dataset, we found significant increases in receptor tyrosine kinase (RTK) and epithelial to mesenchymal transition (EMT) pathways, which may indicate cells increasing this signalling to promote drug resistance^2^.

The SCTK is flexible and integrated well with other Bioconductor tools through the use of a SummarizedExperiment based object. These objects allow the user to transfer the data and results performed within the SCTK to other Bioconductor tools that use the SummarizedExperiment or SingleCellExperiment objects without needing to convert it. In addition, scRNA-Seq data and analysis from other sources can be converted into a object compatible with the SCTK using a single function.

We have developed the Single Cell Toolkit (SCTK), the first fully interactive toolkit that allows a user to perform standard scRNA-Seq workflow interactively in R. With this toolkit users can process data, visualize their results, and save the data into a convenient object for further downstream analysis. Because the SCTK uses the SCTKExperiment object, the resulting data object is compatible with other tools that accept SummarizedExperiment or SingleCellExperiment objects. The toolkit supports various use cases, ranging from data visualization of pre-processed data to a complete scRNA-Seq analysis pipeline from filtering to pathway activity analysis.

The SCTK is available through Bioconductor (https://bioconductor.org/packages/release/bioc/html/singleCellTK.html). Detailed installation instructions are available on the SCTK help website (https://compbiomed.github.io/sctk_docs/). Analysis scripts and detailed walkthroughs from this abstract are available as vignettes in the package.

## Methods

The SCTK is organized into multiple interactive tabs that divide up the scRNA-Seq workflow. Below we describe our example datasets and analysis tools that are available through the interactive SCTK package and GUI.

### Mucosal-associated Invariant T (MAIT) Cells

To demonstrate how interactive analysis can be performed in the SCTK, we will use an example dataset of mucosal-associated invariant T (MAIT) cells^15^. A set of 96 CD8+ MAIT cells were sorted, 47 cells were stimulated with cytokines, and the cells were processed and sequenced using the Fluidigm C1 system. The data was included with the MAST package. Cytokine stimulation of MAIT cells results in increased cytokine gene expression and pathway activity changes that can be identified with the differential expression analysis and pathway activity analysis modules of the SCTK, serving as an effective control for our toolkit methods. The results of this analysis are detailed in **Supplementary Note 1**.

### Pluripotent Stem Cells

A dataset demonstrating batch effects in single cell data was created by Tung, et. al^14^. Three induced pluripotent stem cell lines were sequenced in triplicate on the Fluidigm C1 platform using a total of 9 plates. The resulting data had a clear batch effect that could affect downstream analysis if it is not corrected. After removing the batch effect, the experimental replicates should not separate during analysis, allowing the data to be used to identify biological differences between the individuals. The results of this analysis are detailed in **Supplementary Note 1**.

### Data Upload

After installing the SCTK, users can start the Shiny app by running the `singleCellToolkit()` function with a SCTKExperiment object as an input to automatically load the data into the app. Alternatively, a user can choose to upload a count matrix directly through the Shiny app by uploading a text file, along with optional sample and feature annotation files. The SCTK will create a SCTKExperiment object to store the toolkit analysis results. This object can be exported after analysis has been completed. In addition, experienced computational researchers can run individual functions and analyses available in the SCTK from the command line.

### Data Summary and Filtering

After scRNA-Seq data has been loaded into the SCTK, a table of data summary metrics is available for display to the user. Because scRNA-Seq data is very sparse, dataset specific filtration and normalization can affect downstream analysis^16^. By displaying the number of samples, genes, average number of reads per cell, average number of genes per cell, and the number of genes with low or no information, the user can make decisions about how best to filter their data for downstream analysis. Users can also delete outlier samples, filter the dataset based on an annotation, filter genes by a feature annotation, and modify the annotation information. The filtering applied while using this tab modifies the underlying data that is used throughout the app. A snapshot of the original uploaded data is preserved so a user can always return to the original uploaded data to restart the analysis or try a different filtration protocol.

### Dimensionality Reduction and Clustering

Visualization of scRNA-Seq data is crucial to identifying subclusters of cells present in the data. Dimensionality reduction techniques allow a user to visualize scRNA-Seq data by summarizing the observed variation into lower dimension space. Principal component analysis (PCA) transforms the matrix into components that describe the variation observed in the data. An alternative to PCA, t-distributed stochastic neighbor embedding (t-SNE), is also frequently used when analyzing scRNA-Seq data because it is able to embed a large amount of variation into a small number of dimensions and works well on sparse data^17^. When users open the dimensionality reduction and clustering tab in the SCTK, a list of available reduced dimension datasets and algorithms is provided. Because these algorithms can take a long time to compute on large datasets, users can precompute the reduced dimension data and store it in a SCTKExperiment object before uploading the data into the SCTK. For smaller datasets, users can perform PCA and t-SNE directly through the SCTK app. The resulting reduced matrices will be stored in the underlying object that can be downloaded when analysis is complete. The resulting data can be displayed in the dimensionality reduction and clustering tab. Annotation information can be added to the plot by selecting annotations with which to color or shape the points in the scatterplot.

After visualization of the data, users may want to stratify the scRNA-Seq data into clusters that appear during dimensionality reduction. Users can choose to cluster their data using k-means clustering, hierarchical clustering, or CLARA (Clustering for Large Applications). Clustering is typically performed on the PCA data, because t-SNE data does not retain the distance between clusters in its results. After the clustering algorithm is complete, the plot is automatically updated to display the resulting clusters. If the user wants to save the cluster results, the cluster assignments can be stored in the annotation data frame of the SCTKExperiment object and visualized on other reduced dimension data or used as an annotation in other analysis tabs. Additionally, other clustering algorithms can be run on the command line, saved as annotation information in the SCTKExperiment object, and visualized in this tab.

### ComBat Batch Correction

Because of the complexities of the library preparation and the low starting material in scRNA-Seq experiments, non-biological variation (batch effects) are present and can be a major source of variation present in single cell experiments^18^. ComBat is a widely used method for adjusting for batch effects in microarray and RNA-Seq data^19^. If users identify variation associated with a technical effect, ComBat can be run within the SCTK to remove this variation before further downstream analysis. Users can choose an annotation present in the annotation data frame and add additional covariates to the ComBat model before performing batch correction. After batch correction, the ComBat results are stored as an additional assay in the SCTKExperiment object, which can then be used in the other analysis tabs within the SCTK.

### Differential Expression and Biomarker Creation

Differential expression analysis is used to identify genes that are up or down regulated between conditions. Users can apply differential expression algorithms commonly used for bulk RNA sequence including limma^20^ and DESeq2^21^, or perform an ANOVA to identify differentially expressed genes by selecting one or multiple condition variables present in the annotation information. Users can customize the differential expression results by changing the number of genes to return, the p-value significance cutoff, and the p-value correction method applied to the results. The resulting gene list is displayed as a table and also in a heatmap which can also be customized. Users can download the gene list directly or create a biomarker list for a specific cell type or cell cluster, which can be stored in the gene annotation information in the SCTKExperiment object.

Single cell RNA-Seq specific tools for differential expression have been developed that can accommodate some of the unique characteristics of scRNA-Seq data. MAST, Model-based Analysis of Single-cell Transcriptomics, has been developed to address these issues by using a hurdle model^15^. MAST has been implemented within the SCTK. Users can choose whether to use MAST’s adaptive thresholding model, choose fold change and expression thresholds, and identify significant genes based on conditions present in the annotation information provided. The results are presented in a table, violin plots, or visualized in a heatmap and can be saved as a biomarker in the SCTKExperiment object or downloaded directly.

### Subsampling and Differential Power Analysis

The relative complexity of scRNA-Seq experimental designs makes it difficult for investigators to ensure that an experiment will have sufficient power while operating on a finite budget. Whereas there are tools for optimizing bulk RNA-Seq designs^22,23^, these fail to account for the tiered nature of scRNA-Seq experiments, where each biological replicate may contribute any number of cells to be sequenced, each of which may belong to one of many cell types or subpopulations. Users of the SCTK can project estimated power metrics based on their dataset with variable simulated parameters including sequencing depth, number of sequenced cells, and number of biological replicates. To produce results within a reasonable timespan, the Shiny interface only allows users to vary one parameter at a time while keeping the others fixed. The command line allows users to probe all parameters at once, producing multidimensional power estimates which will help investigators optimize their scRNA-Seq experimental designs.

### Pathway Activity Analysis

Gene expression measurements can be summarized into a signature or set of genes to create a score that represents the activity of that set of genes in a sample. By summarizing genes in known signaling pathways, cells with active signaling pathways or specific cellular functions can be identified. Gene Set Variation Analysis (GSVA) uses gene sets to create these signatures^24^. The molecular signature database (MSigDb) is a database of molecular signatures that can be used in GSVA^25^. GSVA has been implemented in the SCTK. Users can select their input data, gene set(s), and GSVA parameters interactively through the app. After GSVA is complete, scores will be displayed in either violin plots or a heatmap on the Pathway Activity tab of the SCTK. Users can save the pathway activity scores into the annotation data columns of the SCTKExperiment object or download the scores directly.

## Acknowledgements

We thank Zijian Han, Ziyan Li, Zichun Liu, and Shiyi Yang for contributions to the MAST wrapper code. This research was supported by U01CA220413 from the NIH.

## Author Contributions

W.E.J conceived and supervised this research; J.D.C. and M.Y supervised this research; D.F.J, T.F, E.B, S.C.P, and S.C contributed to the software; W.E.J and D.F.J wrote the manuscript.

